# Encoding models uncover fine-grained feature selectivity for bodies, hands and tools

**DOI:** 10.64898/2026.04.09.717525

**Authors:** Davide Cortinovis, Martin N. Hebart, Stefania Bracci

## Abstract

Category-selective areas in the occipitotemporal cortex (OTC) are typically characterized by broad tuning, yet neuroimaging suggests a finer-grained organization reflecting distinct computational roles. We combined image-level fMRI with artificial neural network (ANN)-based encoding models to investigate the selectivity and feature sensitivity of category-selective areas in ventral and lateral OTC. Using densely sampled fMRI data in three participants across six sessions, we identified functional dissociations between body, hand, and tool responses at the individual image level. Area-specific encoding models accurately predicted responses to millions of novel images, maintaining clear category preferences. Importantly, comparisons between models trained on areas selective for the same category revealed distinct feature sensitivities consistent with the areas’ anatomical location and hemispheric lateralization. These findings provide evidence for fine-grained specialization within OTC and demonstrate how ANN-based encoding models can uncover the computational, feature-level basis of category selectivity.

## Introduction

A substantial body of evidence indicates that the ventral temporal cortex contains category selective areas that exhibit preferential activation to specific object categories, such as faces, body parts, or scenes^1^. Recently, artificial neural networks (ANNs) have been adopted to model category selectivity in human visual cortex^2^. In particular, ANNs have been combined with encoding approaches^3,4^ to generate “virtual” models of category-selective areas and to predict their responses to novel images^5,6,7,8,9^. For example, one study^10^ trained encoding models on fMRI data from functionally localized face-, body-, and scene-selective areas; when tested on large sets of novel images, the models showed robust category selectivity, thus validating the functional tuning of these cortical areas.

While ANN-based encoding models have successfully captured broad category selectivity in visual cortex, human neuroimaging reveals finer-grained distinctions in both ventral and lateral occipitotemporal cortex (VOTC and LOTC, respectively^11^). Selectivity is mirrored across both VOTC and LOTC, as both regions show selectivity for faces, bodies, hands, and inanimate objects^12,13,14,15,16,17,18,19^. This selectivity has also been observed with high-field fMRI^20^ and intracranial recordings^21^. Furthermore, category-selective responses for the same object categories (e.g., bodies, hands) often exhibit distinct functional profiles depending on their anatomical location (LOTC vs. VOTC) or hemispheric lateralization. For instance, the ventral and lateral tool-sensitive areas respond to different tool-related properties, with the lateral area more sensitive to shape and action-related properties and the ventral area to surface properties^22^ (see also^23^). These ventral and lateral distinctions have been proposed to reflect partially dissociable computational roles, with ventral regions supporting object recognition and lateral regions supporting action-related properties^24,25^.

Category-selective areas also exhibit hemispheric asymmetries, with stronger right-hemisphere responses for bodies and left-hemisphere responses for hands^12,18^. Finally, with higher spatial resolution or minimal smoothing, even classic category-selective areas can be further parcellated into multiple clusters, each responsive to the same broad category^26,27,28,29^.

For example, multiple limb-selective areas have been identified bilaterally^27^, arranged along a body-part map^30^ or according to action-related principles^22,31^. Converging evidence further shows that the representational content of category-selective areas is itself finer-grained, encoding features beyond canonical category boundaries^32,33,34^.

Considering this evidence, our objectives in the present study were twofold. First, we leverage image-level functional analysis and ANN-based encoding models to provide a stringent test of selectivity for anatomically close and partially overlapping areas. Second, we use the encoding models to explore differential feature representations in areas exhibiting the same category selectivity.

Using densely sampled fMRI data from three participants scanned across six sessions, we localized areas selective for whole bodies, hands, and tools using a functional localizer. Participants then viewed 200 images depicting a range of body parts and inanimate objects, allowing assessment of functional selectivity at the image level^35^. Then, we trained encoding models on these neural responses following the approach of^10^ and tested them on a large set of novel images. To interpret model predictions, we applied occlusion-based saliency mapping to identify image regions that most strongly contributed to predicted neural responses. Finally, beyond validating fine-grained category selectivity for closely overlapping voxels, we used the encoding models to probe whether areas selective for the same category differ systematically in their underlying feature sensitivity, with such differences reflecting the position of each area in ventral or lateral OTC and across hemispheres, in line with the distinct computational goals these regions support.

Our results demonstrate that image-level functional selectivity and encoding models can dissociate body-, hand-, and tool-selective areas in ventral and lateral OTC and reveal that areas labelled as selective for the same category can substantially differ in the features they encode, consistent with the computational role of the region (ventral vs. lateral OTC) or the hemisphere.

## Results

The current work had two main objectives. First, we characterize functional selectivity at the level of individual images in body-, hand-, and tool-selective areas of ventral and lateral OTC. Second, we use ANN-based encoding models to provide stringent tests of category selectivity and to uncover feature-level differences between areas selective for the same category, with a focus on dissociations related to ventral and lateral OTC and to the two hemispheres.

We localized areas selective for whole-bodies, hands, and tools in three participants and recorded event-related responses in those areas to a stimulus set containing 200 images of body-parts (whole-bodies and hands) and inanimate objects (tools, manipulable, and non-manipulable; see Figure 1a). First, we tested the functional selectivity of category-selective areas; then, we trained encoding models based on the activations of those areas and evaluated the functional responses of those models.

**Figure 1.**
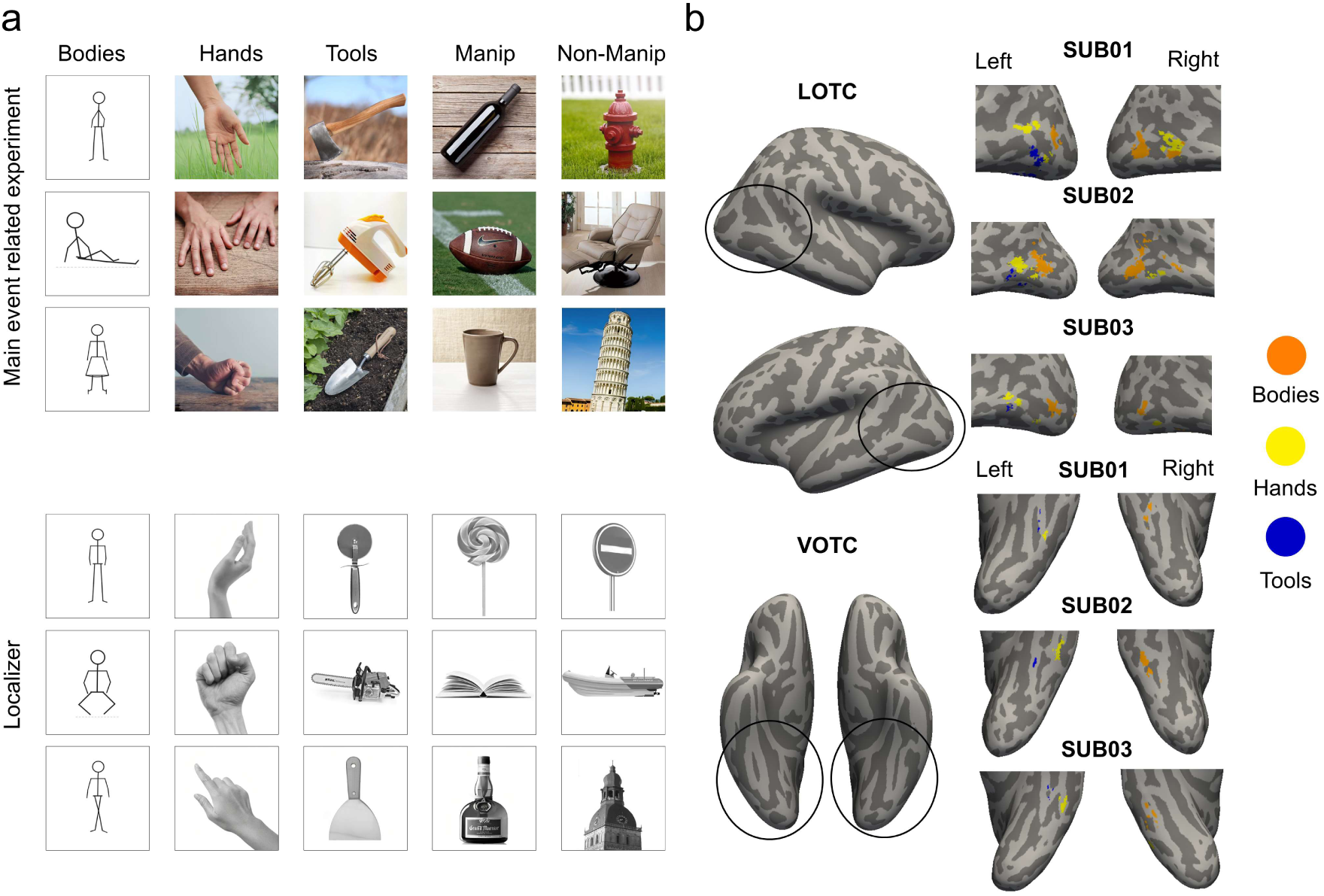
Stimulus sets and ROIs. a) Stimuli used in the main experiment (above) and in the localizer (below). Both sets included the same 5 categories: whole-bodies, hands, tools, manipulable, and non-manipulable objects. b) Single subject category-selective activation maps. Activations for body, hand, and tool selective areas in bilateral ventral and lateral occipitotemporal cortex (p < .0001 uncorrected at the voxel level, FDR < .05 at the cluster level). ROIs were selected in volume space and projected for visualisation on the Freesurfer-reconstructed native surface of each participant^93^. Images of bodies were replaced with stick figures.

**Figure 2.**
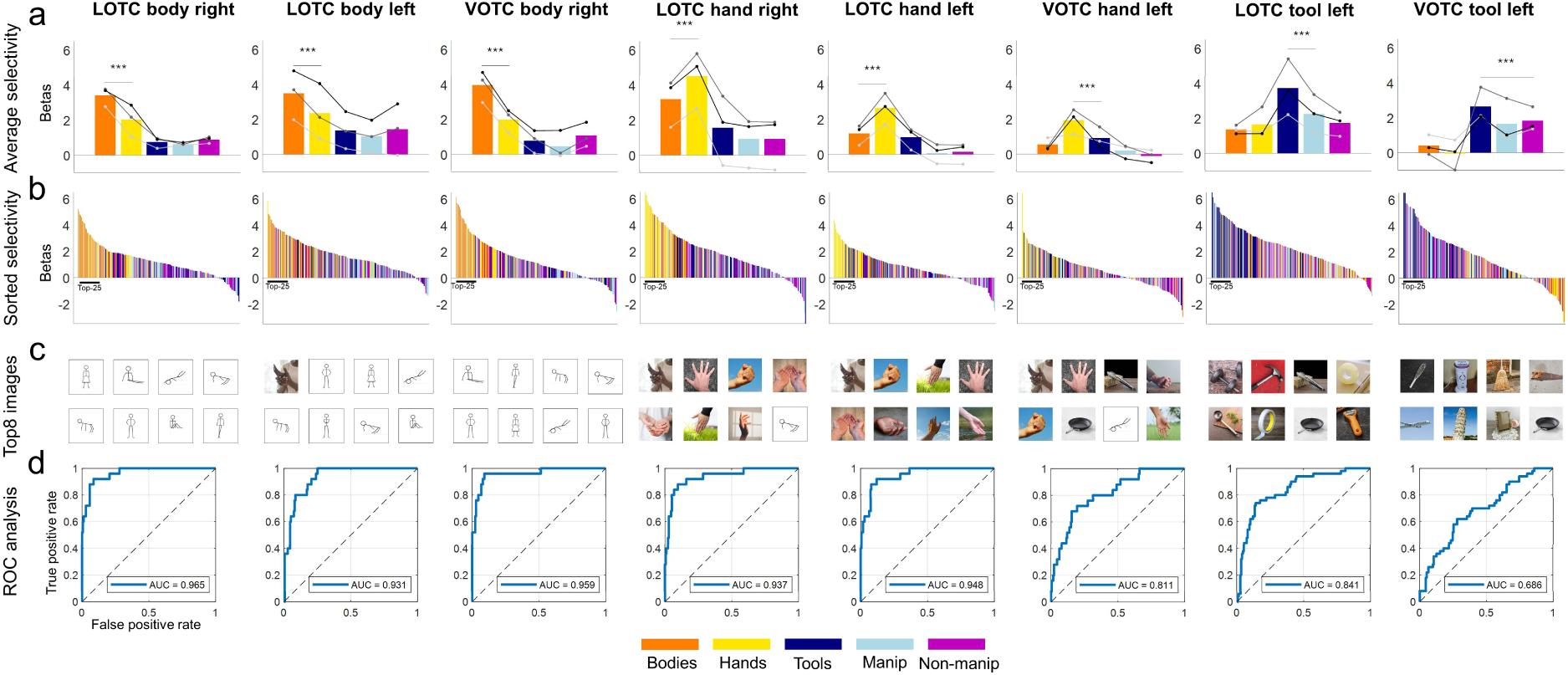
Functional selectivity profiles across neural category-selective areas. a) Average selectivity by category. Lines indicate the response for each single subject. Stars indicate significance (*p* < .001) between the highest activation and the second highest activation. b) Sorted selectivity (from highest to lowest) for each stimulus, averaged across subjects. Black line under each plot indicates the top-25 most activating stimuli. c) Top-8 images activating each area. d) Area under the Receiver Operating Characteristic curve (AUC). The AUC is a ratio of the true positive rate vs. false positive rate, measuring how well neural responses discriminate between the preferred stimulus class from all other stimulus categories. Images of bodies were replaced with stick figures.

### Ventral and lateral OTC areas selective for bodies, hands, and tools

In each participant, we identified areas selective to images of whole-bodies, hands, and tools in ventral and lateral OTC (*FDR* cluster-corrected at *p* < .05) with an independent localizer (see Methods). We only included those areas that could be reliably localized in all three participants (Figure 1b). Whole-bodies activated bilateral regions in LOTC surrounding the lateral occipital sulcus, further extending more anteriorly and superiorly in the right hemisphere; in VOTC, a body-selective area could be reliably identified only in the right hemisphere, around the occipitotemporal sulcus. In LOTC, hands exhibited stronger activations in the left hemisphere, spanning the posterior inferior temporal gyrus and extending more anteriorly and superiorly towards the middle temporal gyrus; albeit smaller, a hand-selective area could also be identified in the right hemisphere; in both cases, hand selectivity was anterior to body selectivity; in VOTC, hand selectivity was identified exclusively in the left hemisphere, around the occipitotemporal sulcus. In LOTC, tools elicited a strongly left-lateralized activation in the inferior temporal gyrus, neighbouring but anterior and inferior to the hand cluster; a smaller cluster of responses was observed in VOTC, around the medial fusiform gyrus. Notice that the ROIs were defined with contrasts that were specifically chosen to exclude voxels that respond to more than one category (hands and tools or hands and bodies).

As a first step, we assessed the internal consistency of the fMRI data in the identified ROIs using a split-half procedure using the main event-related experiment data (see Methods). We found that with the full set of stimulus repetitions, the response patterns demonstrated consistent reliability (LOTC-body right: Spearman-Brown corrected *r* = 0.54; LOTC-body left: *r* = 0.53; VOTC-body right: *r* = 0.48; LOTC-hand right: *r* = 0.58; LOTC-hand left: *r* = 0.58; VOTC-hand left: *r* = 0.3; LOTC-tool left: *r* = 0.48; VOTC-tool left: *r* = 0.38).

Next, we characterized functional selectivity at the level of individual images, allowing us to assess fine-grained distinctions between closely related categories (e.g., bodies vs. hands). The functional selectivity of each ROI is visualized in Figure 3. We quantified selectivity using three primary measures: a *d’* index, a Receiver Operating Characteristic (ROC) analysis, and a top-N analysis. The *d’* index provides a preference score for a category: values around 0 indicate no preference, values above 1 indicate moderate selectivity, and values above 2 indicate strong selectivity. The significance of the *d’* for each ROI’s preferred category was tested against the *d’* of the other non-preferred categories using 10,000 permutations. The ROC analysis assesses how well the neural activation can be used to distinguish one category from all others, giving a classification measure where 1 represents perfect separation (i.e., all stimuli from the preferred category elicits more activation than all the other stimuli) and 0.5 represents chance level. Finally, the top-N analysis determines the proportion of stimuli from an area’s preferred category that were present within the top 25 most activating images.

**Figure 3.**
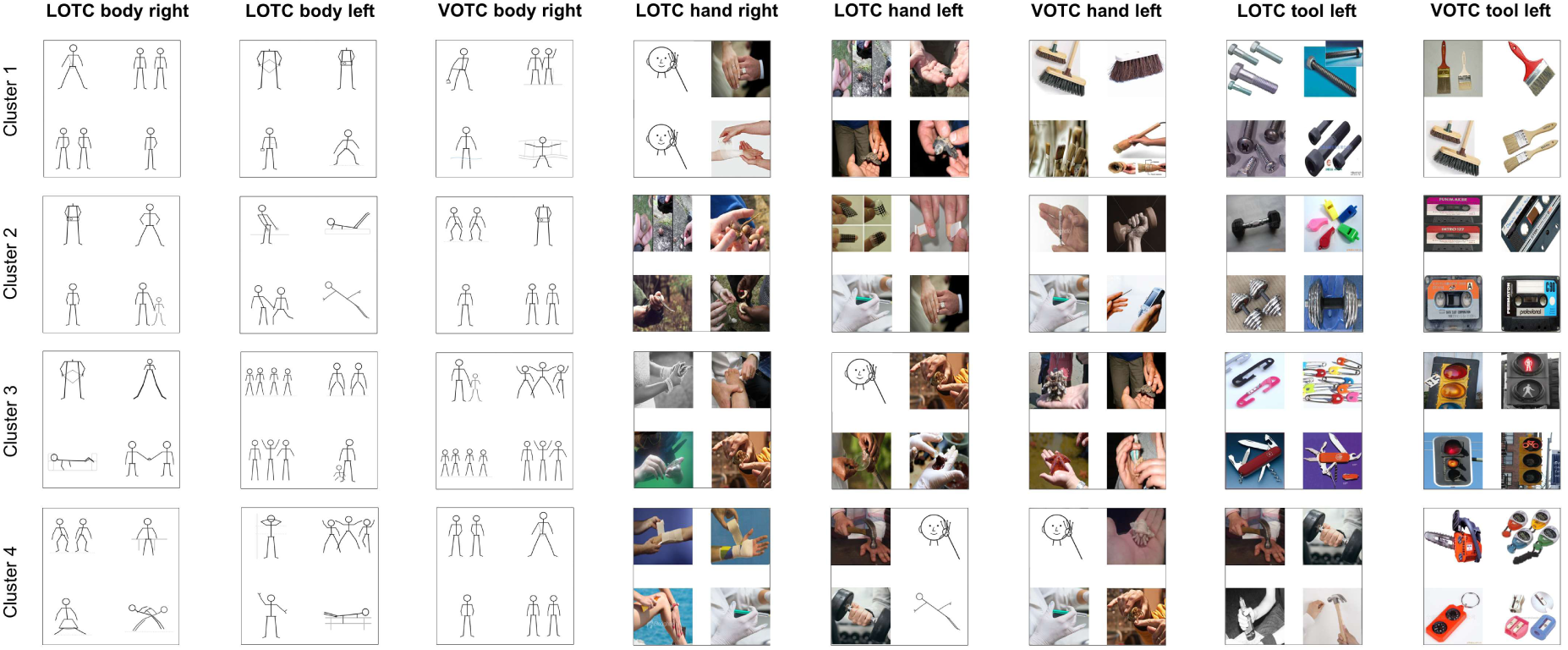
Encoding modelling analysis. Predictions for each image in each encoding model was sorted from highest to lowest, and images sharing similar visual content among the top 2500 images (based on the penultimate layer of a ResNet-50) were clustered. Each cluster was ranked based on the average selectivity of the images contained therein. The top 4 images are shown for each cluster. Images of faces and bodies were replaced with stick figures.

The ROIs showed significant and robust category selectivity. For body-selective areas, LOTC-body right: *d′* = 2.44; *AUC* = 0.97 (indicating a 97% probability that a randomly chosen body stimulus would elicit a higher response than a non-body stimulus), and 72% of *top-25* responses were bodies; LOTC-body left: *d′* = 2.0; *AUC* = 0.93; *top-25* = 64%; VOTC-body right: *d′* = 2.48; *AUC* = 0.96; *top-25* = 76%. For hand-selective areas, LOTC-hand right: *d′* = 2.1; *AUC* = 0.94; *top-25* = 68%; LOTC-hand left: *d′* = 2.3; *AUC* = 0.95; *top-25* = 64%; VOTC-hand left: *d′* = 1.2; *AUC* = 0.81; *top-25* = 44%. Finally, for tool-selective areas: LOTC-tool left: *d′* = 1.38; *AUC* = 0.84; *top-25* = 72%; VOTC-tool left: *d′* = 0.72; *AUC* = 0.69; *top-25* = 52%.

The *d’* for the preferred category was statistically significant (*p* < .0125, Bonferroni corrected with 4 comparisons) against the *d’* for the other categories in all ROIs except in VOTC-tool, which generally showed weaker selectivity and more similar responses between all inanimate object categories (*d’* tool vs manipulable: *p* = 0.013; *d’* tool vs non-manipulable: *p* = 0.044); this weaker selectivity suggests that ventral tool-selective responses may reflect broader inanimate object properties rather than tool identity per se, an interpretation we return to below.

Together, these findings confirm that OTC contains spatially distinct clusters tuned to bodies, hands, and tools. In VOTC, we found right-lateralized body selectivity and left-lateralized hand selectivity but found weak and non-significant tool selectivity relative to other inanimate objects. These results replicate and extend previous parcellations of OTC^12,13,18,20,36^, reinforce hemispheric lateralization for body, hand, and tool responses^13,18^, and confirm that hand and tool selectivity is stronger in lateral than ventral OTC^37^.

### Brain encoding models confirm fine-grained category selectivity

While functional selectivity analysis confirmed the possibility of dissociating body-, hand-, and tool-selective responses in OTC, we sought to investigate the feature spaces underlying category-selective areas, while providing a robust and unbiased test of their selectivity. Following the approach of^10^, we trained a series of encoding models to predict the mean fMRI BOLD response within each ROI from image features extracted from a deep neural network (ResNet-50, see methods for details). After training, we evaluated the performance of the models and used them to perform an in-silico screening of large, independent sets of images (ImageNet^38^; ecoset^39^), to identify the visual features that drove the highest predicted activations, thereby revealing the learned tuning properties of each brain area.

Results show that the encoding models for all category-selective areas were highly successful in predicting their activation patterns. For LOTC-body right the model achieved a prediction accuracy of *r* = 0.67 against a group noise ceiling (reported as √r_sB_) of 0.8; LOTC-body left: *r* = 0.59 (noise ceiling [0.74]); VOTC-body right: *r* = 0.66 (noise ceiling [0.72]); LOTC-hand right: *r* = 0.67 (noise ceiling [0.78]); LOTC-hand left: *r* = 0.66 (noise ceiling [0.78]); VOTC-hand left: *r* = 0.41 (noise ceiling [0.55]); LOTC-tool left: *r* = 0.51 (noise ceiling [0.64]); VOTC-tool left: *r* = 0.45 (noise ceiling [0.56])). Notice that the noise ceiling was computed as a (Spearman-Brown corrected) split-half on the averaged group data (contrary to the reliability analysis, which was performed at the subject level), thus representing an overestimation (upper bound) of the noise ceiling, and indicating high reliability of the data.

Since we were able to successfully train area-specific encoding models with good performance, we then proceeded to evaluate their responses to a separate set of images. Specifically, we screened large image datasets and evaluated the response prediction of each encoding model for all images. For a first qualitative assessment, we visualized the images that were predicted to most strongly activate each model (see Supplementary Material); then, to move beyond idiosyncratic repetitions of nearly identical images, we also applied an unsupervised clustering analysis to the top 0.1% of images (≈2,500). Visual features were extracted from the penultimate layer of a ResNet-50 pretrained on ImageNet, and k-means clustering with cosine distance was used to group images by shared visual characteristics. This approach allowed us to identify the most consistent features across top-ranked stimuli, to provide a more comprehensive picture of each model’s tuning preferences, and to test for the presence of recurrent features that are not related to the hypothesised preferred category. In this section, we describe the general response of body-, hand-, and tool-trained encoding models; in the following sections we target more specifically the distinct feature properties and representations that distinguish the different areas.

A qualitative inspection of the top-ranked images revealed striking and highly specific tuning preferences for each model (see Figure 4 for examples after clustering procedure and Supplementary Materials for the “raw” unclustered top-100 most activating images). For models trained on body-selective areas, the top images consistently depicted whole human bodies, often capturing the full body form rather than isolated body parts. Clustering confirmed this preference: the majority of images within each cluster belonged to human bodies. Similarly, for the models trained on data from hand-selective areas, the top-ranked images (including in each cluster) almost exclusively featured human hands in various postures, both in isolation and interacting with objects. Importantly, the images depict hands and not the general body form, further confirming dissociable responses to hands from other body-parts; additionally, VOTC-hand left seemed to be sensitive to specific types of inanimate objects, such as tools like brooms or brushes. Finally, the model trained on data from LOTC-tool left identified a preference for tools and highly manipulable objects: the top images in each cluster consisted of a diverse array of graspable, functional objects, including nails, weights, silverware, paperclips, and Swiss-army knives, and hands interacting with objects (such as hammers); visualising the overall most activating images also show strong activations for more stereotypical tools such as hammers, pliers, scissors. Despite their varied appearances, the common characteristic was that they were all highly manipulable man-made objects designed for a specific function; all of them were also handheld objects. Many (but not all) objects had a metallic surface and presented an elongated handle. In line with the above functional selectivity analysis, VOTC-tool left presented a different pattern, responding to some tool objects (such as brushes) but also to other inanimate – and often squared – objects in general (stoplights, cassette tapes).

**Figure 4.**
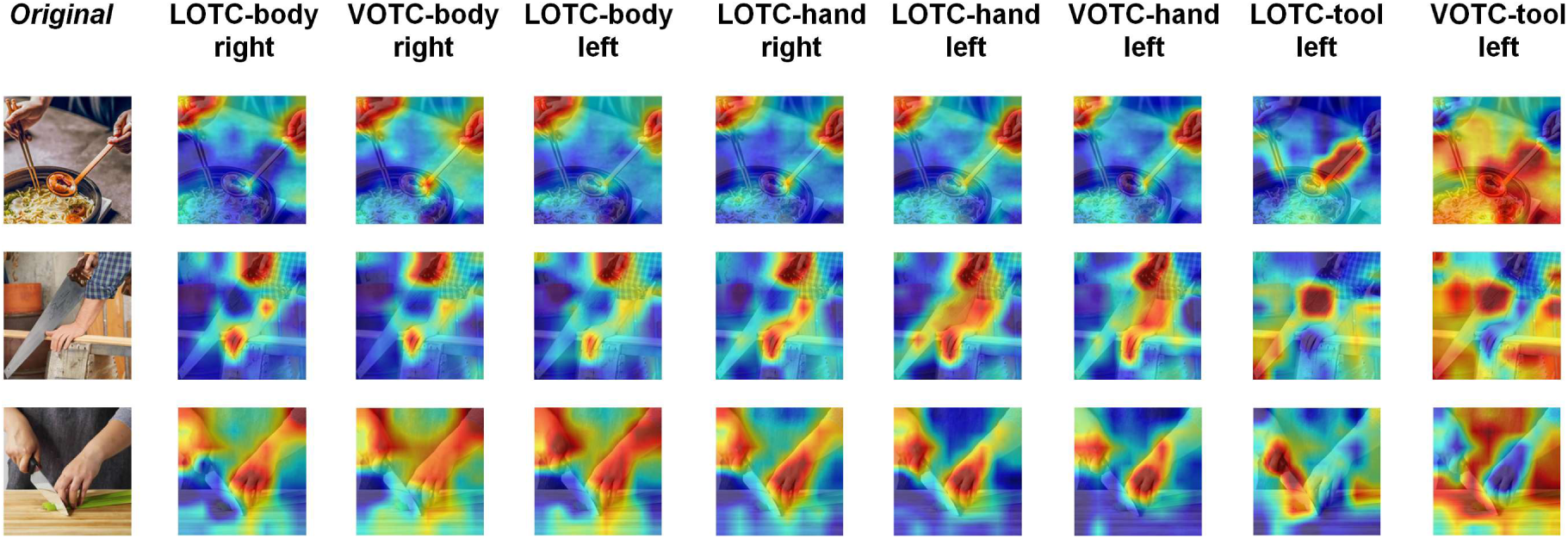
Saliency (“heatmaps”) analysis. 2000 masks (8×8) were generated that randomly occluded parts of the image, and the effect on the encoding models’ predictions were calculated. Areas of the images that elicit a decrease in response are color-coded in red, thus highlighting their saliency. Original images from the COCO dataset included the full body and were thus replaced with images only showing hands and arms.

These findings provide evidence for the expected category selectivity and against the influence of potential stimulus selection bias. Even when challenged with a massive and diverse stimulus set, the encoding models consistently identified images belonging to each area’s preferred category as the most effective stimuli. This suggests that the category selectivity observed in these areas is a genuine and robust property of the visual system.

The tuning to the preferred category in each model was further confirmed by the salience maps analysis (Figure 4). We presented encoding models with images containing the exact same scenes depicting body-parts interacting with objects and tools, sampled from the COCO dataset^40^. Each encoding model generally revealed preference within the image for their respective category: body models responded to parts of the images depicting body-parts such as arms and shoulders (sometimes including – but not specific to – hands), and at the same time showing no preference for inanimate objects; hand models were specifically tuned to hands (and generally not other body-parts); and tool models preferred highly-manipulable objects such as scissors, brushes, or cutlery, with the encoding model trained on VOTC-tool data exhibiting a more general preference for inanimate objects.

### Distinct feature sensitivity underlies areas selective for the same category of objects

The analyses conducted so far provide evidence for category-specific processing of objects within each selective area. However, even areas that respond to the same category may process distinct underlying features. For example, tools are often both elongated and metallic; two regions might respond similarly to a tool, while representing different aspects of the stimulus (e.g., shape versus surface properties). To investigate such potential feature sensitivities, we compared the prediction scores of encoding models across areas. Specifically, we computed pairwise differences between the models’ predictions for each image and identified the images that maximally activated one model relative to another. We performed these contrasts both across hemispheres (e.g., LOTC-hand right vs. LOTC-hand left) and across ventral and lateral OTC (e.g., LOTC-hand left vs. VOTC-hand left). This analysis revealed systematic and interpretable feature-level dissociations between areas selective for the same category, aligning with known distinctions between ventral and lateral OTC and hemispheres. Results can be visualized in Figure 5, where we show representative images for each contrast.

**Figure 5.**
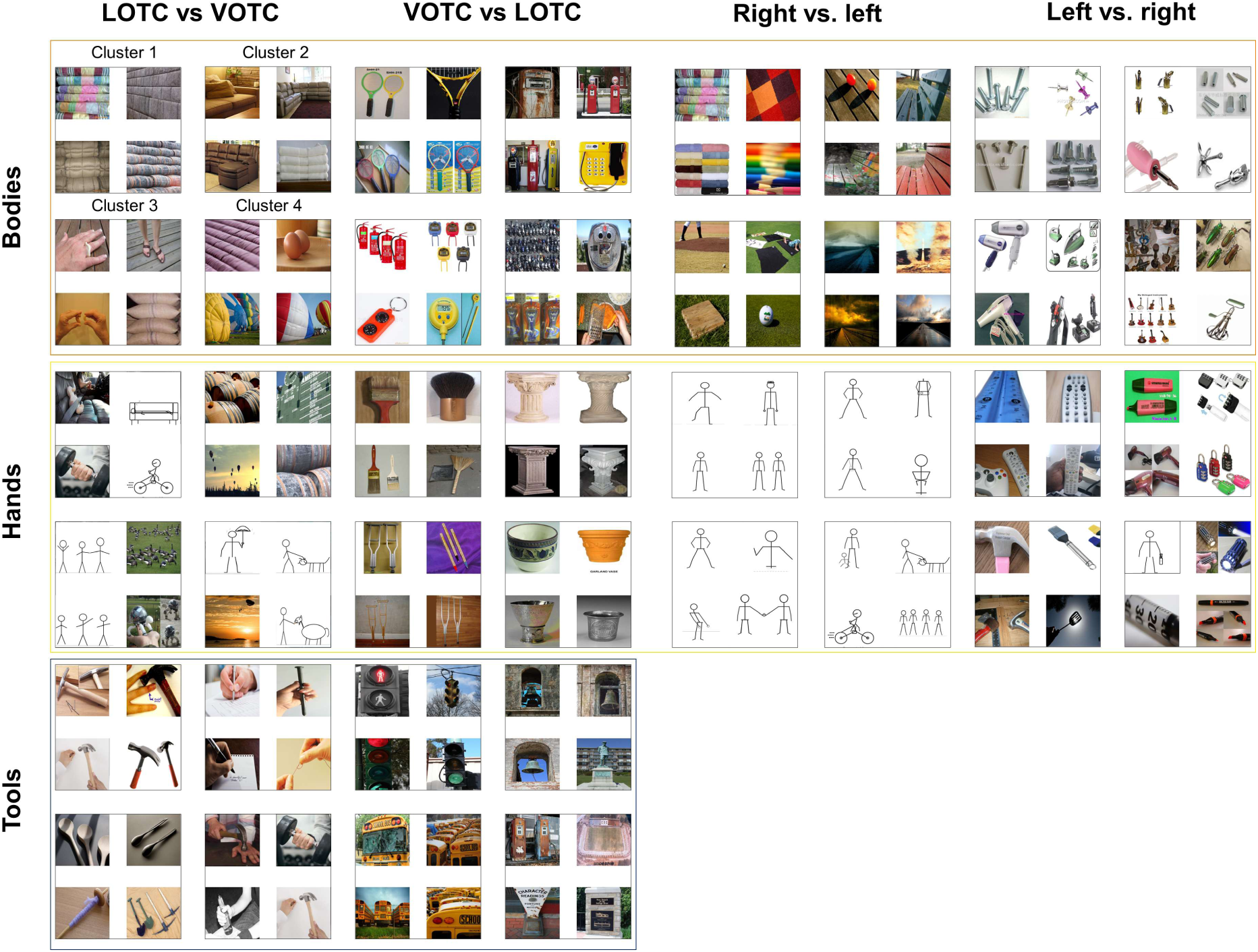
Distinct feature preferences across areas responding to the same category. Encoding models trained on different areas selective for the same category were compared by pairwise subtracting their predictions scores. The top images were then ranked and clustered. Each cluster was ranked based on average selectivity. Example images are shown for each cluster. Images of faces and bodies were replaced with stick figures.

When contrasting hand-selective areas in the two hemispheres, we found that LOTC-hand left responded strongly (relative to LOTC-hand right) to handheld manipulable objects, typically tools such as hammers, markers, remote controls, and spatulas. By contrast, LOTC-hand right showed a clear preference (relative to LOTC-hand left) for images of bodies, often embedded in richly textured backgrounds. These results make sense in light of their hemispheric lateralization: as shown by previous studies, the left hand-selective area neighbours and partially overlaps with tool selectivity and to a less extent to manipulable objects^12,14,22^, whereas the hand-selective area, in the right hemisphere, neighbours body-selectivity^41^, which usually forms a more extended cluster in the right hemisphere (at least in right-handers^42^). Interestingly, LOTC-hand left did not simply show stronger tool responses than VOTC-hand left. Instead, LOTC-hand left was driven by more complex scenes containing people interacting with animals or multiple objects arranged in rectilinear (e.g., buildings) or curvilinear (e.g., clustered umbrellas) layouts, whereas VOTC-hand left was more responsive to single elongated objects against simple backgrounds (e.g., columns, brooms on a uniform-coloured background). The fact that sensitivity to tools cannot distinguish between ventral and lateral hand-selective areas is again consistent with the action-related gradient: tool selectivity is “sandwiched” between lateral and ventral hand-selective areas, and VOTC-hand has also been implicated in representing action-related object properties^22^. The observed differences may therefore reflect distinct sensitivities to low- or mid-level visual features such as elongation and curvilinearity, properties that have been found to explain large portions of OTC in general and hand and tool selectivity in particular (e.g.^43,44^), or may underlie sensitivity to complex scenes involving interactions, consistent with the recently proposed interaction sensitive lateral pathway^25,45^.

Comparisons between ventral and lateral tool-selective areas revealed a similar division of labor. LOTC-tool left was preferentially driven by classically defined tools and manipulable objects, often depicted with hands (e.g., hammers, pens, syringes). In contrast, VOTC-tool left responded more broadly to inanimate objects, particularly large, non-manipulable items such as streetlights, buildings, vehicles, and bell towers. These results support a division between lateral tool-selective regions tuned to action-related objects and ventral regions tuned more generally to large non-graspable inanimate objects^46,47^.

Finally, differences between body-selective areas across the two hemispheres were less straightforward to interpret. Contrasting models trained on body-selective areas across hemispheres revealed that – perhaps surprisingly – differences were driven primarily by mid-level visual features and material properties. Images driving LOTC-body left more strongly than LOTC-body right included inanimate objects, such as tennis rackets, fire extinguishers, phones, and small metallic objects, such as nails; conversely, images that strongly activated LOTC-body right, but not LOTC-body left were characterized by saturated colours, rectilinear and repetitive textures, and materials such as textiles, fabrics, and wood. A similar distinction emerged when comparing lateral and ventral regions in the right hemisphere: LOTC-body right responded more strongly to images containing rectilinear features, including some form of textiles, fabrics, wooden pavements, and, additionally, sofas; VOTC-body right was preferentially driven by small circular or stubby objects, such as flyswatters, fire extinguishers, or small phone booths. The relative difficulty of interpreting these contrasts likely reflects greater feature similarity across body-selective areas compared to hand- and tool-selective areas. Indeed, the average activation difference between LOTC-tool left and VOTC-tool left (≈ 3.6) and between LOTC-hand left and LOTC-hand right (≈ 2.3) was notably larger than that between body areas (≈ 1.7). In other words, the more similar two models’ prediction scores are, the less distinctive, and thus less interpretable, the feature differences that separate them become.

Together, these results demonstrate that body-, hand-, and tool-selective areas exhibit robust image-level functional selectivity, while ANN-based encoding models provide a stringent test of such selectivity. Areas selective for the same category differ systematically in the features they encode, with these differences aligning with known distinctions between ventral and lateral OTC and between hemispheres^22,25^.

## Discussion

In this study, we first characterized functional selectivity for category-selective areas in OTC for bodies, hands, and tools at the level of individual images, and then trained ANN-based encoding models on those responses. These models captured fine-grained category preferences observed in the brain and revealed distinct feature representations underlying selectivity for areas responding to the same category. We show that body-, hand-, and tool-selective areas are reliably dissociable across ventral and lateral OTC and exhibit both strong selectivity for their preferred category and graded responses to non-preferred categories based on their hypothesised computational goal. Encoding models trained on each area’s responses predicted neural activity with high accuracy, and large-scale image screening confirmed that the strongest predicted activations consistently belonged to the expected category. Crucially, comparisons between encoding models revealed systematic differences in feature sensitivity among areas selective for the same category, such as heightened sensitivity to action-related properties in the lateral tool-selective area and more general processing of the features belonging to many inanimate objects in the ventral “tool” area. Together, these findings indicate that category-selective areas in OTC reflect multiple, partially dissociable computational roles that depend on their position along ventral vs. lateral OTC or their hemispheric lateralization.

Research has consistently shown that discrete regions in OTC respond selectively to specific object categories^1^, including faces, bodies, scenes, words^41,48,49,50^, and more recently food^51,52,53^. However, converging evidence suggests that this organization is more fine-grained than this canonical map implies. Areas traditionally treated as unitary, such as the Fusiform Face Area or the Extrastriate Body Area can be parcellated into multiple clusters with distinct functional profiles^18,27,36,54^. For body-parts specifically, previous work has identified limb-selective patches arranged in a ring-like fashion around motion-sensitive cortex^27,28^, as well as dissociable responses to hands versus whole bodies^13,30,31^.

Despite this evidence, much of the literature still treats large regions such as EBA as homogeneous units, averaging across heterogeneous subareas. This practice risks obscuring the complexity of cortical organization and underestimates the degree to which OTC responses are distributed across distinct, specialized clusters. Importantly, such distinctions are not a trivial, secondary phenomena: hand-selective voxels, for example, support specific functions distinct from body-selective voxels, including sensitivity to hand pose and connections to parieto-frontal circuits for tool use^12,55,56,57^, and even have distinct developmental trajectories^58^. Recognizing this finer-grained organization is essential for understanding how OTC supports distinct feature spaces and behavioral goals^59,60^.

Our study advances this debate by providing a robust test of functional selectivity for body-, hand-, and tool-selective areas. At the image level, these areas showed distinct and graded response profiles: hand-selective areas responded preferentially to hands over whole bodies and exhibited strong secondary responses to tools; tool-selective areas responded most strongly to tools and highly manipulable objects rather than to all small inanimate objects. This interpretation is consistent with a graded organization extending from effector-specific representations within hand-selective areas and the hand–tool overlap^12,14^ to more general processing of manipulability in more ventral and anterior regions of lateral OTC^22,37^.

Beyond investigating OTC functional responses, our work builds on and extends ANN-based encoding approaches used to model responses in ventral visual cortex^5,7,9,10,61,62,63,64,65,66^. Following the rationale of^10^, we trained encoding models on the mean response of category-selective areas and tested their functional profiles on large-scale image datasets. Encoding models are particularly important given the inherent constraints of fMRI and the limited sampling of image space achievable even in large-scale neural datasets (e.g.^67,68,69,70^). By serving as virtual instantiations of category-selective areas, these models provide a complementary approach to these datasets: once trained, they can be evaluated on millions of images, enabling fine-grained tests of selectivity and representational structure that would be impractical with direct human experiments alone^10^.

Here, we successfully trained encoding models based on the responses of body-, hand-, and tool-selective areas, enabling interpretation and visualization of the visual features that differentiate both across category-selective areas and within areas responding to the same category. Whereas body-selective areas showed relatively homogeneous sensitivity to body images, hand- and tool-selective areas could be dissociated along an action-related dimension. Left hemisphere hand-selective areas were more sensitive to end-effector objects (e.g., tools such as hammers or scissors), while right hemisphere hand-selective areas showed greater sensitivity to the body form. A similar dissociation emerged within tool-selective cortex: ventral regions showed reduced sensitivity to action-related features and were more broadly tuned to inanimate object properties, whereas lateral regions were more selectively tuned to canonical tools and end-effectors objects. Importantly, these findings demonstrate that encoding models are not limited to testing predefined hypotheses about category selectivity: by probing the functional space of trained models, it is possible to identify the visual features that drive regional selectivity, thereby moving beyond “word models” of category organization^10,71^. This approach enables principled differentiation between areas that respond to the same category but encode distinct feature dimensions (see also^61^). In line with this, hemispheric differences within hand-selective cortex were captured by differential sensitivity to action-related objects, with left-lateralized tuning for inanimate objects carrying high action information and right hemisphere regions preferentially encoding body-related features. This pattern is consistent with the left-lateralization of the hand–tool overlap^12^ and previously reported action-related topographic organization^22^.

Moreover, the encoding models allowed us to differentiate areas in the ventral and lateral pathways that are sensitive to tools. While the lateral tool area is more tuned to properties related to high manipulability, the ventral tool area is not selective to tools per se but is biased towards processing inanimate objects in general, including large objects (see^15^ for a different perspective on tool “selectivity” within the medial fusiform gyrus). Additional visual properties such as elongation and metallic appearance also contributed to model predictions, in line with recent fine-grained characterizations of responses to manipulable objects^72^; however, we think that these features, while important, are not the primary features represented within lateral tool areas, that might be instead involved in representing action-related properties of objects. Notice that the features that are mapped via the encoding models are extracted from a CNN, that might be biased towards visual properties and be unable to access the action-related properties characterising tools (see^22^); thus, shape and material properties may be just a proxy for the action-related features represented by the tool (and hand) areas. Overall, these results are consistent with the distinct processing between lateral and ventral OTC, one involved in representing action-related properties (and shape), and one possibly dedicated to the elaboration of surface properties of objects, that are typical not of tools specifically but of many object categories^37,73^.

While our results support the existence of distinct category-selective clusters, they also intersect with a complementary view in which OTC is organized along shared representational dimensions such as animacy^74,75,76^, real-world size^77^, aspect-ratio^74^, and action^22^. Recent work suggests that neighbouring and partly overlapping activations for whole-bodies, hands, tools, manipulable objects, and even food in lateral OTC^12,13,72,78^ can be explained by action-related dimensions describing how body parts interact with objects^22^. Our findings do not contradict this perspective: while our aim was to dissociate as much as possible functional responses using targeted contrasts to maximise selectivity, we also observe that each selective area presented a graded response to the non-preferred categories. For instance, while hand-selective areas are most strongly driven by hands, the images that next-best activated the area were body-parts (specifically when legs were prominent) and tools, consistent with an action-related representational gradient, where hands and tools share similar effector properties. Thus, a category-based account coexists with a dimensional framework^32,33^, and ANN-based encoding models offer a powerful way to bridge the two. Therefore, our results are consistent with a more graded view of object preference^35,79^. Finally, these results open avenues for directly comparing biological and artificial representations in models that capture both functional and spatial organization of visual cortex^80,81,82,83^, providing insights into the computational pressures shaping cortical topography^84,85^.

In summary, our study both confirms and extends classic accounts of category selectivity in OTC. It confirms them by demonstrating robust selectivity for bodies, hands, and tools, and extends them by revealing systematic feature-level differences within and across category-selective areas, offering a computational framework for understanding how functional specialization emerges in OTC. When provided with this feature information, encoding models can reliably distinguish between whole bodies, hands, and tools, categories that are often grouped together or not considered. Even if not ascribing to a strict modular view of category selectivity, our analyses confirm the presence of information related to hands and tools in visual cortex, suggesting that it is important to consider these categories or the domain they support to have a complete picture of the ventral visual cortex organization and topography.

## Methods

### Participants

Three right-handed participants (2 females, age range 24-30 years) with normal or corrected-to-normal vision and no history of neurological disorders took part in the study. Participants provided informed consent, and the experimental procedures were approved by the Ethics Committee of the University of Trento.

### Stimuli

Two distinct sets of stimuli were used for the main event-related experiment and the localizer. The main experimental stimulus set consisted of 200 coloured images with natural backgrounds, primarily sourced from the THINGS dataset^86^ and internet searches. The images belonged to 5 categories: 25 whole bodies, 25 hands, 50 tools, 50 manipulable objects, and 50 non-manipulable objects. Tools were defined as objects that are used with hands to physically interact with other objects or surfaces (e.g., scissors, knives); manipulable objects were graspable but are usually the passive receiver of the action (e.g., books, cups); non-manipulable objects are large non-graspable objects (e.g., vehicles, buildings).

The localizer stimulus set consisted of 72 greyscale images divided into 6 categories: whole bodies, hands, tools, manipulable objects, non-manipulable objects, and chairs. Each category included 12 images presented on a white background, adapted from a set used by^87^.

Inanimate object categories were controlled for visual differences like shape and orientation. All images were resized to 400×400 pixels and subtended 8° of visual angle.

### Experimental Design and Procedure

Each participant completed six scanning sessions over two weeks, totalling approximately nine hours of fMRI data per participant. Data for both experiments were acquired in each session. Visual stimuli were presented using the Psychophysics Toolbox^88^ in MATLAB (2021b, The Mathworks), projected onto a screen and viewed through a mirror mounted on the head coil.

#### Main Experiment

The main experiment used a rapid event-related design. In each run, lasting 5 min and 24 sec, 100 images drawn from the full stimulus set were presented for 500 ms with a 3500 ms inter-stimulus interval (ISI). Blank trials were included approximately 20% of the time. Participants performed a catch trial detection task, pressing a button whenever an image of a bug or plant appeared (∼10% of the time). Around 60 runs per participant were collected, totalling on average 15 repetitions per image.

#### Localizer

The localizer followed a block design. In each of 6 runs, lasting 5 min 36 sec, blocks of images from a single category were presented. Within a block, images were shown for 400 ms with a 266 ms ISI. 5 block repetitions per category were included. Participants performed a one-back task, pressing a button when an image was repeated twice in a row (one repetition per block).

### MRI Acquisition and preprocessing

All imaging data were collected at the Center for Mind/Brain Sciences (CIMeC), University of Trento, using a 3T Siemens Prisma scanner with a 64-channel head coil. Functional volumes were acquired for both the localizer and main experiments using a T2*-weighted echo-planar imaging (EPI) sequence with a multiband acceleration factor of 3 (parameters: TR = 2,000 ms; TE = 28 ms; flip angle = 75°; 69 axial slices covering the whole brain; FoV = 220 mm, voxel size of 2 x 2 x 2 mm). A high-resolution T1-weighted anatomical image was also acquired in the first scanning session for each participant using an MPRAGE sequence with a voxel size of 1 x 1 x 1 mm.

Preprocessing was performed using the Statistical Parametric Mapping software (SPM12; Wellcome Trust Centre for Neuroimaging) in MATLAB (R2021b). The pipeline included slice-timing correction, head motion correction via spatial realignment to the first volume of each run, and coregistration of the functional data to the participant’s anatomical scan. To maximize spatial precision and avoid possible overlap between neighbouring category-selective voxels, the functional images were analyzed in the native participant space. A FWHM Gaussian kernel of 3 mm was applied to the localizer data to improve signal-to-noise ratio and increase localization’s robustness. Runs with head motion exceeding predefined thresholds (2 mm in translation or 1 mm in rotation) were excluded from the analysis.

A general linear model (GLM) was then fitted to the preprocessed data. For the main experiment, each stimulus presentation was modelled as a unique event. For the localizer, the six object categories were modelled as conditions. In both cases, the six motion-correction parameters were included as nuisance regressors. Predictors were generated by convolving a boxcar function with SPM’s canonical hemodynamic response function.

### Region of Interest (ROI) Selection

Category-selective regions of interest (ROIs) were defined on the native cortical volume of each participant using data from the localizer experiment. Analyses were focused on the ventral and lateral occipitotemporal cortex (VOTC and LOTC respectively). Functional ROIs were identified using individualized contrasts for category, corrected for multiple comparisons at a cluster-level threshold (*FDR* < .05). Areas selective for bodies were identified with a contrast of whole-bodies vs. hands and tools; areas selective for hands were identified with a contrast of hands vs. whole-bodies and tools; areas selective for tools were identified with a contrast of tools vs. all other categories. The contrasts were specifically chosen to functionally dissociate these highly related and often overlapping representations. Indeed, in lateral OTC, voxels responding to bodies, hands and tools partially overlap along a continuum from dorsal posterior to ventral anterior^22^; here, by employing direct contrasts we specifically selected non-overlapping voxels responding to one of the three categories.

### Functional selectivity analysis

As a first step, we evaluated the selectivity of the identified ROIs. Our design allowed us to test the functional responses of category-selective areas at the image level with a relatively high number of images. We quantified these category preferences using a *d’* index^35^, defined

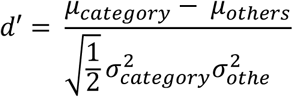

Where *μ_category_* and *μ_others_* are the mean responses to two different categories, and σ_category_^2^ and σ_others_^2^ are their variances. The significance of each category *d’* (vs. the *d’* of each of the other category) was assessed using a permutation test with 10,000 permutations, and all reported results were significant at *p* < .0125 (Bonferroni corrected with *N* = 4 comparisons, each category *d’* vs the other categories *d’*), unless we report no effect. In addition, to provide a complementary assessment of category discriminability, we conducted a Receiver Operating Characteristic (ROC) analysis^35^. For each category, we performed a one-vs-all comparison, treating the stimuli for one category as the “positive” class and all stimuli from the remaining four categories as the “negative” class. The activation values for all 200 stimuli were used as the ranking variable to plot the true positive rate against the false positive rate across all possible thresholds. The Area Under the Curve (AUC) was then calculated as the primary metric of classification performance. An AUC value of 1.0 indicates perfect classification, where all stimuli from the target category elicit a higher response than any stimulus from the other categories, while an AUC of 0.5 represents performance at chance level. Finally, functional selectivity was further tested with a Top-N rank analysis, which indicates the proportion of stimuli from each category among the top 25 responses (e.g., how many hands there are in the 25 stimuli eliciting the highest activation).

### Encoding models

We adopted an analysis framework similar to that of^10^. The encoding model operates by extracting visual features from images using a deep convolutional neural network and learning a mapping from these features to brain activity. This allowed us to predict neural responses to novel images based on their visual content. Then, screening the trained encoding models on large-scale image datasets allowed us to perform a stringent test of category selectivity.

Specifically, we first extracted features for the 200 images using a ResNet-50 architecture pre-trained on ImageNet. The network was chosen based on results from previous studies employing similar encoding modelling procedures^7,10^. We evaluated each ResNet-50 layer and selected the one yielding the highest correspondence between predicted and actual fMRI responses. The best-performing layer was the final global average pooling layer, which outputs a 2048-dimensional feature vector per image. These vectors served as predictors, while the target variable was the average fMRI response to each image across the three participants. The relative low number of images (compared to large scale image datasets, for example the Natural Scenes Dataset^67^) – and hence the possible limits in generalization that can be obtained with this dataset – are due to a trade-off between the number of stimuli and the number of repetitions that can be presented and the quality that can be achieved at an image-level with fMRI.

We used ridge regression to learn a linear mapping from features to fMRI responses and employed 10-fold cross-validation for model evaluation. The dataset was split into 10 disjoint subsets; in each fold, 9 subsets were used for training and 1 for testing. This procedure was repeated so each subset served once as the test set. Within each fold, the optimal ridge regularization parameter (*λ*) was selected through a nested 10-fold cross-validation on the training data to avoid overfitting. Model performance was quantified as the Pearson correlation coefficient (*r*) between predicted and actual responses in each test fold. Final prediction accuracy was calculated as the average correlation across all folds. We trained a separate encoding model for each ROI.

### Reliability of fMRI Response Patterns

To evaluate the internal consistency of the stimulus-evoked fMRI response patterns within each ROI, we conducted a split-half reliability analysis^10,89^. For this procedure, the full set of trials for each of the 200 stimuli was randomly partitioned into two equal halves. The responses for each stimulus were then averaged across the trials within their respective half and across all voxels in the ROI. This process was performed for each participant, and the resulting response patterns were then averaged at the group level, yielding two independent group-average response vectors (one for each split). The Spearman rank correlation was calculated between these two vectors. To estimate the reliability of the full dataset, this correlation coefficient was corrected using the Spearman-Brown (SB) prophecy formula. Because this score represents the maximum explainable variance rather than the maximum achievable correlation, we report the noise ceiling in correlation units as 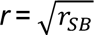 when comparing against model–data correlations^90^. To ensure a stable estimate, this entire process (from random splitting to the corrected correlation) was iterated 100 times, and the final reliability score was taken as the mean of these iterations. To calculate the noise ceiling for the averaged group data, we repeated the same procedure (split-half analysis and Spearman-Brown correction) on the betas averaged across participants. With the entire number of repetitions per stimulus, we obtained reliable fMRI responses (see results). This score functions as a noise ceiling against which we can compare the model performance^89^.

### Image Screening and Datasets

To assess the feature preferences and category selectivity of the encoding models, we screened them against ImageNet^38^ (*n* = 1,281,149), a large-scale image dataset containing diverse object categories traditionally used in machine learning to train models, and ecoset^39^ (*n* = 1,444,892), an ecologically motivated dataset designed to more closely mirror the human visual diet. This combination ensured a broad coverage of categories, including images of people, objects (and specifically, tools and manipulable objects), and their interactions, which are often underrepresented in well-known vision neuroscience datasets such as the Natural Scenes Dataset^67^ (see^91^).

Using the trained models, we predicted BOLD responses for all images in each dataset. Images were then ranked by predicted activation for each ROI, allowing us to identify the categories and features most strongly associated with neural responses. For a first qualitative assessment of category tuning, we first visualize the top activating images ranked by predicted activation for each model; these visualizations enable direct inspection of the visual features and categories that maximally activated each region. However, since the model tends to respond similarly to images that differ only slightly (e.g., multiple near-identical screwdrivers), the top-ranked images can be dominated by a single object or feature. To obtain a more comprehensive picture of the object categories and features underlying the encoding models responses, we therefore applied an unsupervised clustering analysis to the top 0.1% of images (≈2,500) ranked by predicted activation. Specifically, visual features were extracted from these images using the penultimate layer of a ResNet-50 model pretrained on ImageNet. K-means clustering was performed with a cosine distance metric. We visualized the most activating images contained in each of four most selective clusters and generated inferences based on their common visual features. The aim of this analysis was to test if any object category or specific features not related to the hypothesised preferred category elicit strong and consistent activations in the encoding models.

### Interpreting Encoding Model Predictions with Occlusion-Based Saliency Mapping

To understand which parts of the images were driving the model’s predictions, we used an occlusion-based saliency mapping technique inspired by Randomized Input Sampling for Explanation (RISE^92^). This method identifies pixel regions within the image that most contribute to the predicted activation. For this analysis, we selected 25 images from the COCO dataset^40^. All images depicted scenes that included body-parts (such as visible trunks, arms, legs, etc.), hands, and tools (i.e., hairdryers, brushes, scissors, cutlery). The images were randomly sampled from the COCO dataset by selecting relevant labels indicating categories of interest (e.g., “person”, “scissors”), resized and squared to 400×400 pixels. We selected only images in which body-parts, hands, and tools were simultaneously present and sufficiently large to be reliably captured by the masking procedure.

For each image, 2,000 random binary masks (8×8 resolution) were generated and multiplied with the input image to create partially occluded versions. These were passed through the full encoding pipeline (ResNet-50 + ridge regression), and each mask was weighted by the predicted BOLD signal. The final importance map was computed as the weighted sum of these masks, highlighting image regions that most strongly influenced the model’s prediction. This process was applied separately to each encoding model. This analysis was used to gain insight into the image features driving model predictions and to compare feature sensitivity across areas selective for the same category.

### Comparing Category-Selective Encoding Models

Beyond evaluating models in isolation, we also examined pairwise differences between encoding models to uncover feature-level distinctions among brain areas selective for the same category, focusing on contrasts between ventral and lateral pathways and between hemispheres. For each model pair, we identified images that elicited the largest differences in predicted activation, i.e., stimuli predicted to strongly activate one model but not the other. Such differential predictions provide insights into the unique feature sensitivities of each area and offer evidence for potential functional subdivisions underlying feature specificity (and, thus, their computational goals) within category-selective areas. The analysis was conducted on the same large-scale stimulus set used for image screening (ImageNet and ecoset combined, N = ≈ 2,500,000 after removing images common between the two datasets). We focused on contrasts between areas selective for the same category but located either in opposite hemispheres or along different processing pathways. For example, we compared predictions between LOTC-hand left and LOTC-hand right (hemispheric comparison), as well as between LOTC-hand left and VOTC-hand left (pathway comparison).

All predictions were z-score normalized prior to comparison. We adopted the same visualisations and clustering approach as described above: we first plotted the top-activating images with the largest positive differences for each pair of areas, highlighting stimuli that preferentially activated one model over the other; then, we applied the unsupervised clustering analysis to the top 0.1% of images, and plot the images contained in each of four cluster, ranked by their average activations. All analyses were conducted using custom MATLAB scripts.

## Competing interests

The authors declare no competing interests.

## Data and code availability

MATLAB code used for all analyses and the encoding models are available on the Open Science Framework at the following link: https://osf.io/7r9tv

## Supplementary Information

**Supplementary Figure S4.1.**
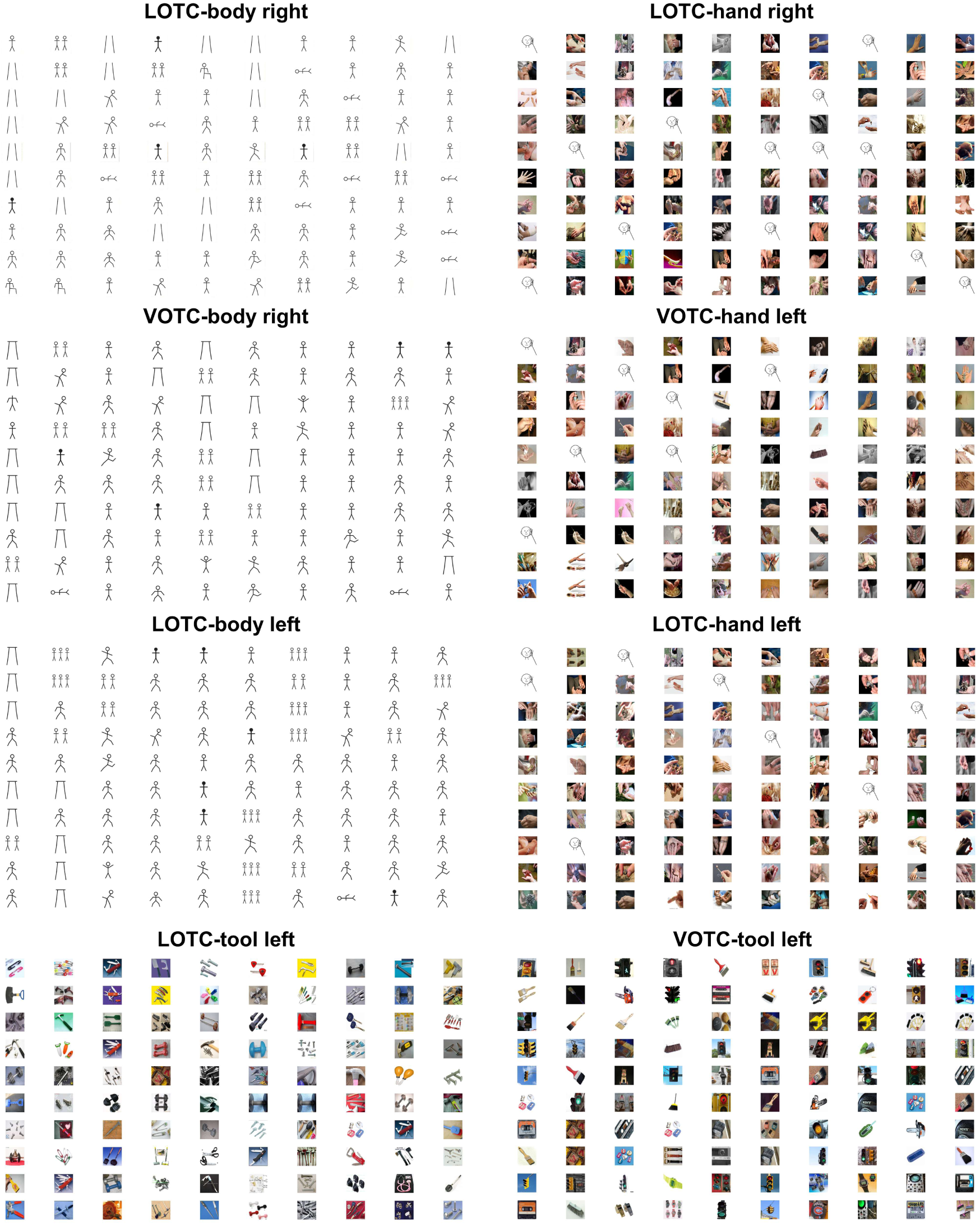
Top-100 most activating images for the body-, hand-, and tool-trained encoding models. Figures of full bodies or hands interacting with faces were replaced with stick figures. Vertical lines in body areas indicate that only the lower body was visible.

## Notes

https://osf.io/7r9tv/

### Competing Interest Statement

The authors have declared no competing interest.

